# Bi-phasic effect of gelatin in myogenesis and skeletal muscle regeneration

**DOI:** 10.1101/2021.05.26.445744

**Authors:** Xiaoling Liu, Er Zu, Xinyu Chang, Ziqi Wang, Xiangru Li, Qing Yu, Ken-ichiro Kamei, Toshihiko Hayashi, Kazunori Mizuno, Shunji Hattori, Hitomi Fujisaki, Takashi Ikejima, Dan Ohtan Wang

**Affiliations:** Wuya College of Innovation, Shenyang Pharmaceutical University, Shenyang, 110016, China; School of Traditional Chinese Materia Medica, Shenyang Pharmaceutical University, Shenyang110016, People’s Republic of China; School of Life Science and Biopharmaceutic, Shenyang Pharmaceutical University, Shenyang110016, People’s Republic of China; Institute for Integrated Cell-Material Science (iCeMS), Kyoto University, Yoshida-Honmachi, Sakyo-ku, Kyoto, 606-850, Japan; Department of Chemistry and Life Science, School of Advance Engineering, Kogakuin University, 2665-1, Nakanomachi, Hachioji, Tokyo, 192-0015, Japan; Nippi Research Institute of Biomatrix, Toride, Ibaraki 302-0017, Japan; Key Laboratory of Computational Chemistry-Based Natural Antitumor Drug Research & Development, Shenyang Pharmaceutical University, Shenyang 110016, Liaoning, P.R. China; Center for Biosystems Dynamics Research (BDR), RIKEN, 2-2-3 Minatojima-minamimachi, Chuo-ku, Kobe, Hyogo 650-0047, Japan

**Keywords:** gelatin, skeletal regeneration, hormesis, ROS, NOX2, IL-6/TNFα

## Abstract

Skeletal muscle regeneration requires extracellular matrix (ECM) remodeling including an acute and transient breakdown of collagen that produces gelatin. However, the physiological function of such a remodeling process on muscle tissue repair is unclear. Here we elaborate on a bi-phasic effect of gelatin in skeletal muscle regeneration, mediated by hormetic effects of reactive oxygen species (ROS). Low-dose gelatin stimulates ROS production from NADPH oxidase 2 (NOX2) and simultaneously upregulates antioxidant system for cellular defense, reminiscent of the adaptive compensatory process during mild stress. This response triggers the release of myokine IL-6 which stimulates myogenesis and facilitates muscle regeneration. By contrast, high-dose gelatin stimulates ROS overproduction from NOX2 and mitochondrial chain complex, and ROS accumulation by suppressing antioxidant system, triggering release of TNFα, which inhibits myogenesis and regeneration. Our findings reveal gelatin-ROS-IL-6/TNFα signaling cascades underlying a hormetic response of myogenic cells to gelatin.

## Introduction

Skeletal muscle has a remarkable capacity to regenerate after mechanical or disease-related injuries. This process often requires changes in biological behavior of myoblasts, the myogenic progenitor cells, located within muscle ECM [1, 2]. Upon damage, quiescent satellite cells (SCs) are rapidly activated to generate Pax7^+^/MyoD^+^ myoblasts that proliferate, migrate into injury section, further differentiate into Pax7^+^/MyoG^+^ myotube and fuse with each other or damaged myofibers to produce new multinucleated myofibers. The myogenesis process is under control of myogenic regulatory factors (MRFs), such as Myf5, MyoD, myogenin (MyoG) that direct myoblasts to enter myogenesis programs and further fuse with myofibers to repair muscle function. Myogenesis process is also signaled and aided by ECM remodeling, growth factors and myokines. The regenerative capability of skeletal muscle is life-long, but nonetheless susceptible to aging, neuromuscular diseases and ECM deficits [3]. However, by what mechanisms ECM regulates the functional, morphological, and molecular events of skeletal muscle regeneration remain poorly understood.

Skeletal muscle is encapsulated by a well-organized network of connective tissue, comprising of ECM with embedded myoblasts, satellite cells and other cell types [4]. The muscle ECM is subdivided into three discrete and interconnected parts, including endomysium (around muscle fiber), perimysium (around bundles of muscle fibers), and epimysium (around the whole muscle) [5]. Collagen (I, III, IV, XII and XIV) is abundant in muscle ECM, accounting for nearly 10% of muscle dry weight. Fibrillar collagen I, the main component of ECM, distributes widely in endomysium, perimysium and epimysium, not only exerting biomechanical support but also influences the biological function of muscle cells [6, 7, 8]. Although it is known *in vitro* that collagen I promotes myogenesis of C2C12 myoblasts [9], muscle regeneration *in vivo* is known to be accompanied by a temporary and local breakdown of collagen, producing a mixture of collagen polypeptides without triple-helicity [10].

During ECM remodeling, fibrillar collagen is digested by collagenase or degraded under inflammation or thermal condition into gelatin that could be further hydrolyzed into smaller peptides by gelatinases such as metalloproteinase-2 and −9 [11, 12, 13]. Under physiological conditions, the amount of gelatin at any given time is balanced by the synthesis and degradation thus maintained at a moderate level [14]. However, a disruption of the synthesis and degradation of gelatin can occur under pathological conditions such as skin inflammation, burn or fever, causing accumulation of gelatin in local tissue and further downstream effects [15, 16]. It was reported that increased generation of gelatin during skin burn activates pro-inflammation cascades and leads to immune dysfunction. Previous studies have shown protective effects of gelatin to reduce TNFα-induced cytotoxicity in murine fibrosarcoma L929 cells, and stimulating effects of gelatin to release pro-inflammatory cytokines by lymphocyte U937 cells [17, 18, 19]; however, it is unknown whether gelatin has influences on myoblasts during muscle repair. In recent years, gelatin as a biomaterial has been found to have wide applications in food industry and tissue engineering due to its excellent biocompatibility, biodegradability and low cost [20]. The gelatin-based biomaterials thus hold great potentials in muscle tissue engineering. *Ostrovidov* et al. fabricated gelatin multi-walled carbon nanotubes to scaffold myotube formation and improve myotube contractions [21], and synthesized gelatin-polyaniline nanofibers to enhance the maturation of excitation-contraction coupling system in myocytes [22]. Gelatin-genipin-based hydrogels drive myogenic cell differentiation thus can potentially be applied to skeletal muscle repair [23]. Although gelatin-based biomaterials are applied widely in tissue engineering including muscle, its biological function on myogenesis or muscle regeneration are unknown.

Gelatin has been previously shown to induce reactive oxygen species (ROS) in multiple cell types to regulate various cell functions. ROS are major mediators of ECM remodeling during regeneration or ECM-related diseases. ROS and antioxidants are associated with skeletal muscle physiology and pathology, suggesting that gelatin may regulate muscle function through ROS signal. ROS in skeletal muscle primarily functions in redox signaling through mechanisms of hormesis, where low-level exposure to ROS elicits beneficial stress adaptation, whereas too high ROS production relative to the antioxidant capacity promotes oxidative stress and cytotoxicity. Interestingly, ROS signal has been reported to both enhance and inhibit myogenesis, depending on its concentration and location. The moderate production of ROS during exercise or regeneration induces myogenic differentiation of satellite cells and myoblasts while excessive accumulation of ROS results in their senescence, apoptosis and regenerative failure in muscle repair [24]. ROS are likely to function as a double-edged sword on myogenesis. It has been reported that the function of ROS on myogenesis is mediated by inflammatory factors, NF-κB p65, FOXO1 [25]. However, the cell and molecular mechanisms underlying the dual effects of ROS on myogenesis are not fully understood.

Herein we have investigated gelatin-induced biological effects on muscle repair and intracellular signaling pathways in C2C12 cells, primary myoblasts, and tibialis anterior (TA) muscle, to elucidate cellular mechanisms underlying the function role of gelatin during myogenesis and muscle regeneration. Our results have revealed a bi-phasic role of gelatin in regulating skeletal muscle repair mediated by intracellular ROS, antioxidant system, and cytokines (IL-6 and TNFα) signaling cascades.

## Results

### Bi-phasic role of gelatin in skeletal muscle regeneration in vivo

To investigate the function of gelatin on skeletal muscle repair, we injected low (LCG, 20 μl of 5 mg/ml) and high (HCG, 20 μl of 20 mg/ml) concentrations of gelatin (saline as control) into CTX-damaged TA muscle 2 days after injury (Fig. 1A). Recovery was evaluated at 7 and 14 d post-injury (D.P.I) by weighing muscle mass and histological examination. Compared to saline, LCG injected mice recovered significantly more TA muscle mass within 14 days, albeit not to the uninjured level (Supplementary Fig. 1A-B). Remarkably, HCG not only did not facilitate recovery, but aggravated original injury, resulting in further muscle loss (Fig. 1B). H&E and IHC staining revealed that at 7 D.P.I, LCG mice formed a higher number of new muscle fibers, characterized by more centrally located nuclei than control mice; whereas little muscle fiber regeneration was observed in HCG mice. At 14 D.P.I., LCG mice displayed tightly packed, well-formed muscle fibers resembling uninjured muscle in PBS mice, but HCG mice had smaller muscle fibers with single nuclei, thus showing impeded or repressed progress compared to the naturally recovered mice (Fig. 1C). We quantified cross-sectional area (CSA) of muscle fibers, number, and percent of myofibers containing centralized nuclei, which consistently showed facilitated recovery in LCG mice and impeded recovery in HCG mice compared to saline group, indicating a bi-phasic effect that at lower concentrations gelatin injection can aid muscle regeneration, but at higher concentrations, it can obstruct muscle regeneration (Fig. 1D-E and Supplementary Fig. 1C-D). We further stained embryonic myosin heavy chain (eMyHC), a skeletal muscle-specific contractile protein expressed during muscle development at 7 D.P.I. Co-staining of eMyHC and laminin revealed that the number of eMyHC^+^/laminin^+^ myofibers was significantly increased by LCG, but reduced by HCG (Fig. 1F-G). This result further supported that gelatin can facilitate muscle regeneration in a bi-phasic and concentration-dependent manner.

**Fig. 1.**
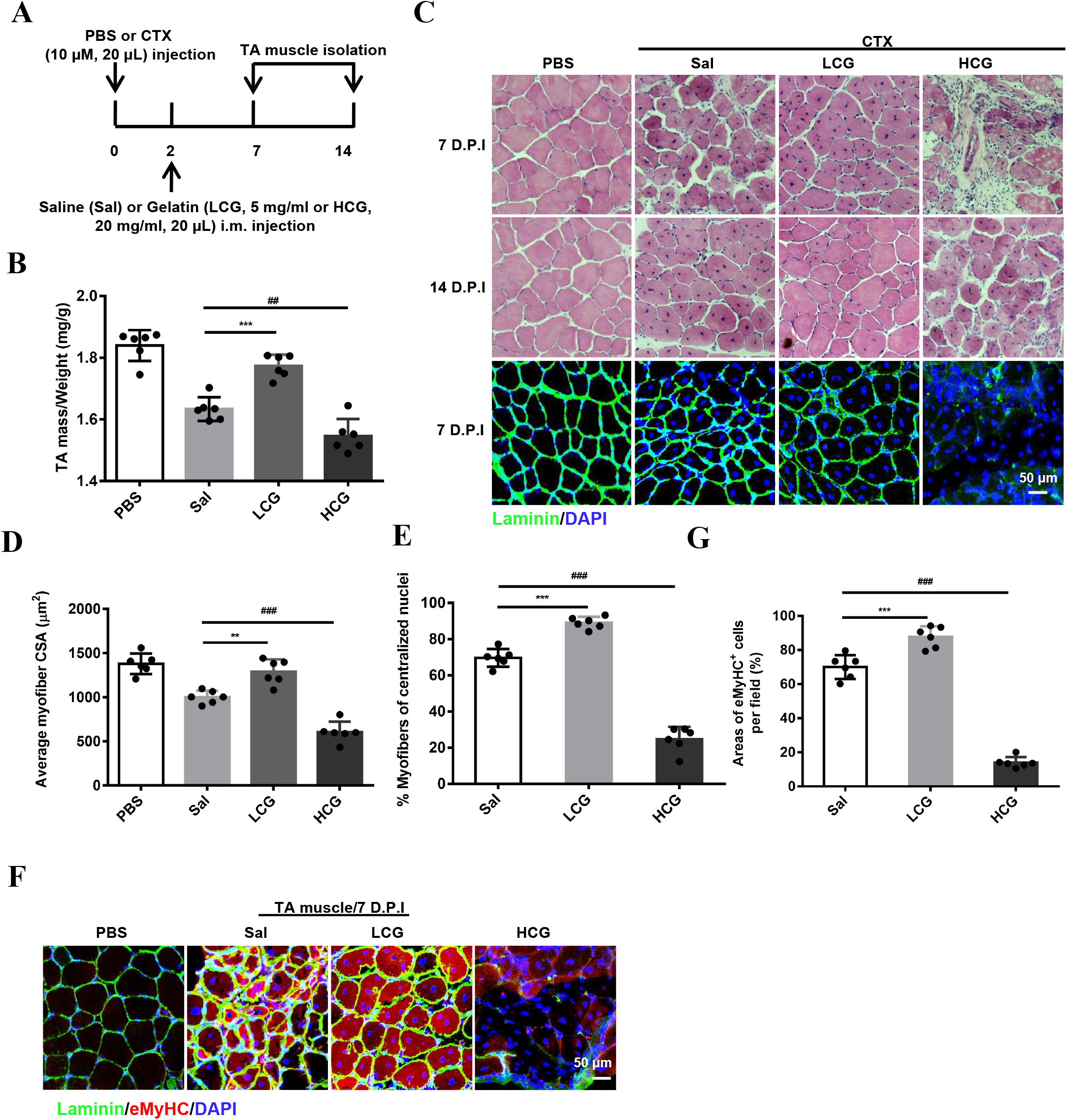
Bi-phasic effect of gelatin in muscle regeneration in vivo. **A.** Experimental schematic diagram depicting CTX injection to induce muscle damage (10 μM, 20 μL in PBS) and gelatin injection (LCG, 5 mg/ml; HCG, 20 mg/ml, diluted with saline) two days post-injury (2 D.P.I.). The TA muscles were collected at 7 and 14 D.P.I. **B.** TA mass relative to animal body mass. **C.** H&E staining of the damaged TA muscle sections at 7 (top row) and 14 (mid row) D.P.I and fluorescence immuno-staining of laminin (green) and DAPI (blue) at 7 (bottom row) D.P.I (n=6). Scale bar, 200 μm. **D.** Average cross-sectional areas (CSA) of regenerated myofibers (n=6). **E.** The percentage of myofibers containing centralized nuclei (n=6). **F.** The fluorescence confocal images of myofibers (green:laminin; red: eMyHC; blue: DAPI) in damaged TA muscle sections at 7 D.P.I (n=6). **G.** The percentage of eMyHC^+^ myofibers area per filed is shown on the right. Significance was determined by unpaired two-tailed Student’s t-test analysis. ^*^*P* < 0.05; ^**^*P* < 0.01; ^***^*P* < 0.001; NS, not significant. Data are means ± SEM.

### Low-dose gelatin promotes satellite cell expansion and the myogenic differentiation

Activation of satellite cells (SCs) and myogenic differentiation are key steps in skeletal muscle repair. Upon damage, muscle satellite cells promptly become activated and start to proliferate, differentiate into myoblasts that migrate, and further differentiate and fuse into multinucleated myofibers. Co-staining of Pax7 and F-actin in TA muscle showed more expansion of SCs in LCG mice but less expansion in HCG mice at 7 D.P.I. (Fig. 2A-B). The results of eMyHC staining in Fig. 1 indicated that gelatin can influence myogenic differentiation. To confirm, we performed IHC staining of MyoG and laminin. The results revealed that LCG extended but HCG reduced the number of MyoG^+^ cells in laminin^+^ cells in the damaged muscle sites (Fig. 2C-D). Consistently with the immuno-staining results, western-blot also showed up- and down-regulated expressions of MyoD and MyoG by LCG and HCG, respectively (Fig. 2E-F). Taken together, during *in vivo* TA muscle regeneration, injecting gelatin into the injury site can promote SCs activation, myoblast differentiation, and fusion, but only when injected at a lower amount such as 20 μl of 5mg/ml.

**Fig. 2.**
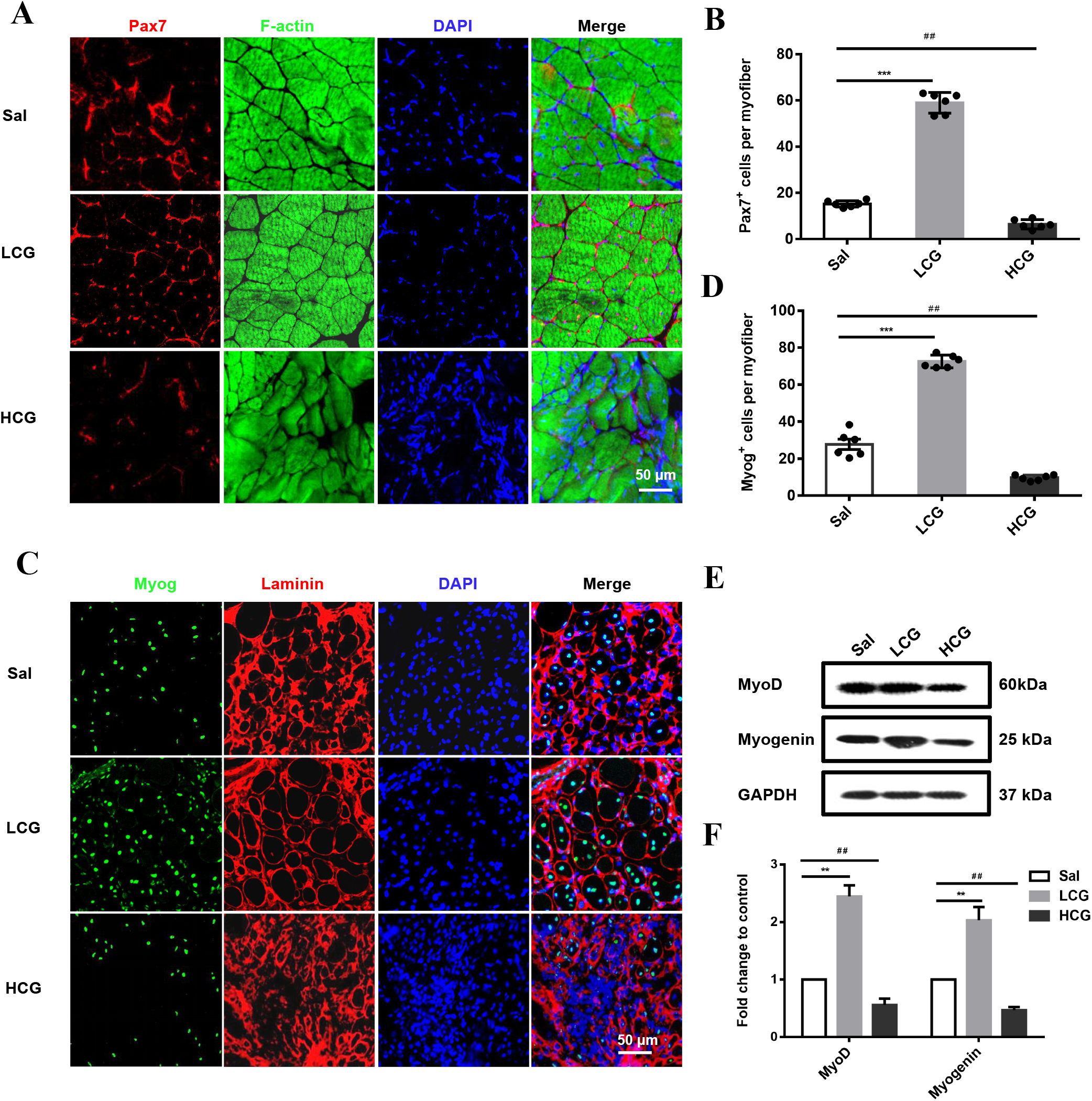
Hormetic effect of gelatin in activating quiescent SC and myogenesis during muscle regeneration. **A-B.** IHC analysis of Pax7^+^ myofibers (red) in TA muscle stained by Alexa Fluor 488 Phalloidin labelled F-actin (green) at 7 D.P.I (n=6). The percentage of Pax7^+^ myofibers area per filed is shown on the right (B). **C-D.** IHC analysis of MyoG^+^ myofibers (green) in damaged TA muscle stained by laminin (red) at 7 D.P.I (n=6). The percentage of MyoG^+^ myofibers area per filed is shown on the right (D). **E-F.** Western-blots of MyoD and Myogenin in damaged TA muscles at 14 D.P.I (n=6). GAPDH is used as a loading control. Significance was determined by unpaired two-tailed Student’s t-test analysis. ^##, **^*P* < 0.01; ^***^*P* < 0.001. Data are means ± SEM.

### Gelatin influences myoblast behaviors in C2C12 myoblasts

To elucidate cellular mechanisms underlying the bi-phasic effect of gelatin, we tested proliferation, migration, and myogenic differentiation of C2C12 myoblasts cultured on gelatin coated dishes (0, 5, 10, 20 mg/ml). Proliferation rate was measured using EdU labeling. After 24 h incubation, cell numbers showed bell-shaped response to gelatin, topped at 5 mg/ml (LCG) and bottomed at 20 mg/ml (HCG) (Fig. 3A-B). Compared to the cells cultured on non-coated dishes or gelatin-coated at 10 mg/ml with large and flattened shape, the cells cultured on LCG appear extended, bipolar and fibroblast-like, while the cells cultured on HCG appear irregular, multi-polar and satellite-like (Supplementary Fig. 2A). Furthermore, the migration of cells is enhanced by nearly 2-fold on LCG dishes, but it is inhibited on HCG dishes (Fig. 3C-F). For myogenic differentiation, LCG not only increased the protein and mRNA expressions of myogenic factors including MyoD, Myogenin, as well as myosin heavy chain (MyHC) (Supplementary Fig. 2B-D), but also elevated the number of MyHC-positive cells (myogenesis index) and promoted the formation of myotubes (fusion index) (Fig. 3G-I) characterized by a wider diameter, a longer length compared with control cells (Supplementary Fig. 2E-F). By contrast, HCG decreased the myogenic differentiation of cells (Fig. 3G-I). Thus, the bi-phasic effect of gelatin on myogenesis *in vivo* is reproduced in C2C12 myoblast cells *in vitro*.

**Fig. 3.**
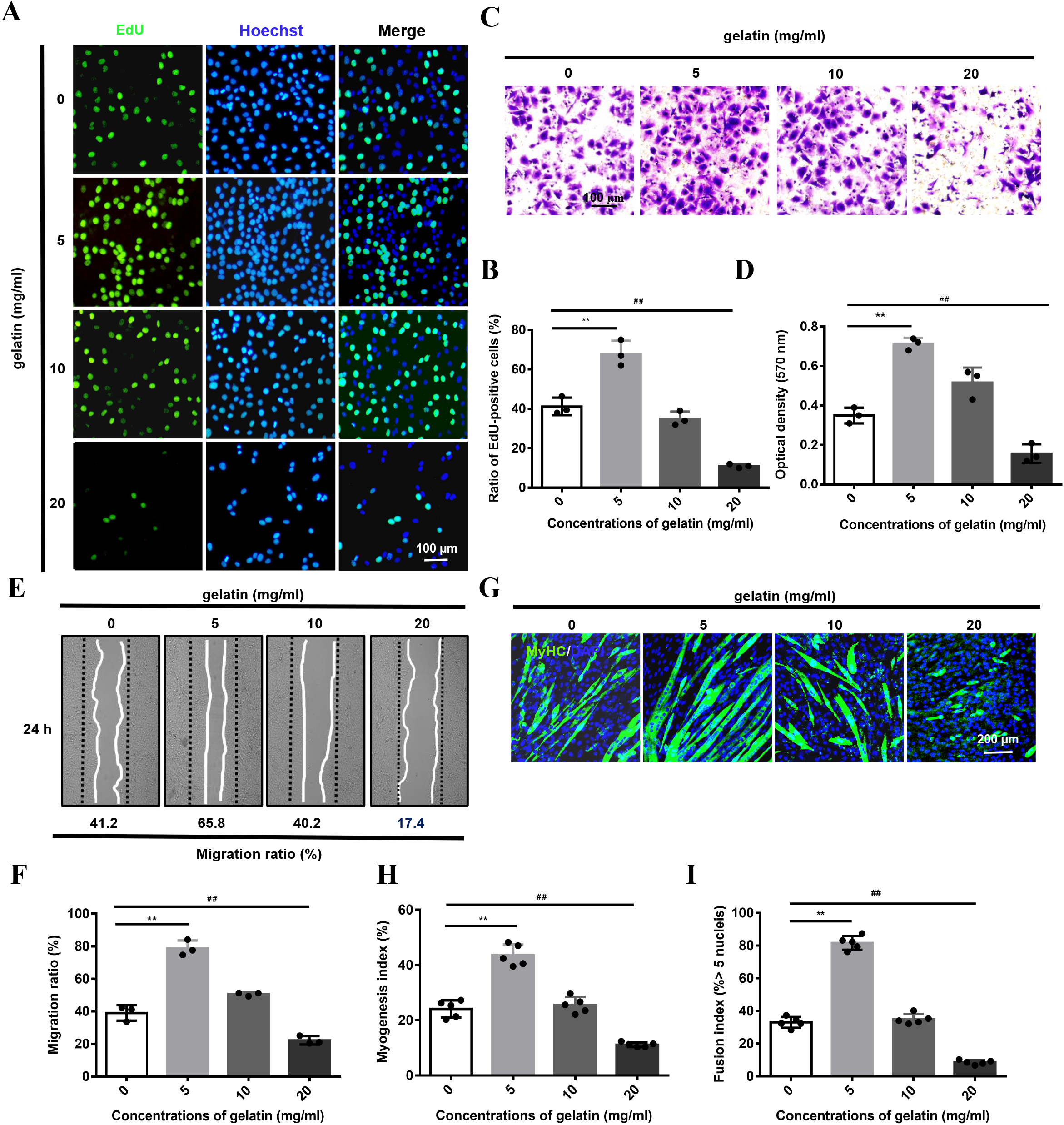
Bell-shaped concentration-response of myogensis to gelatin substrate. **A.** Confocal images of C2C12 on gelatin-coated dishes, stained with Hoechst (blue) and EdU (green). Scale bar, 100 μm. **B.** Right, quantification of EdU-positive (proliferating) cells. **C-F.** Transwell assay (C) and wound healing assay (E) for cell migration measurement and the migration ratio was calculated using Image-Pro Plus software (D, F). **G.** Confocal images of myogenic differentiation of cells stained with an anti-MyHC antibody. Scale bar, 200 μm. **H.** Myogenesis indexes determined as the percentage of MyHC-positive nuclei of the total nuclei. **I.** Myotube fusion index determined as the distribution of the nucleus number in total myotubes. Significance was determined by unpaired two-tailed Student’s t-test analysis. ^##, **^*P* < 0.01. n = 3. Data are means ± SEM.

### Gelatin enhances ROS generation in a dose-dependent manner

ROS have been shown to function as a double-edged sword in multiple cell functions and downstream cellular signaling of gelatin [26, 27, 28]. Using DCFH-DA staining (an indicator for ROS), we found that gelatin stimulated ROS production in C2C12 in a dose-dependent manner (Fig. 4A-B). The cellular concentrations of ROS species such as superoxide radical (O_2_^.−^), hydroxyl radical (·OH), and hydrogen peroxide (H_2_O_2_) all responded to gelatin-coating. LCG significantly enhanced the production of O_2_^.−^; HCG dramatically increased not only O_2_^.−^ but also ·OH (Fig. 4C). We further tested activity of the antioxidative system consisting of antioxidases, including superoxide dismutase (SOD), glutathione peroxidase (GSH-PX), and catalase together with non-enzymatic antioxidants [29]. The results indicated that the activity of antioxidases, especially SOD, was increased by LCG, but gradually inhibited following the increasing concentrations of gelatin. HCG significantly suppressed the activity of GSH-PX and SOD, explaining the aggregative accumulation of ROS (Fig. 4D).

**Fig. 4.**
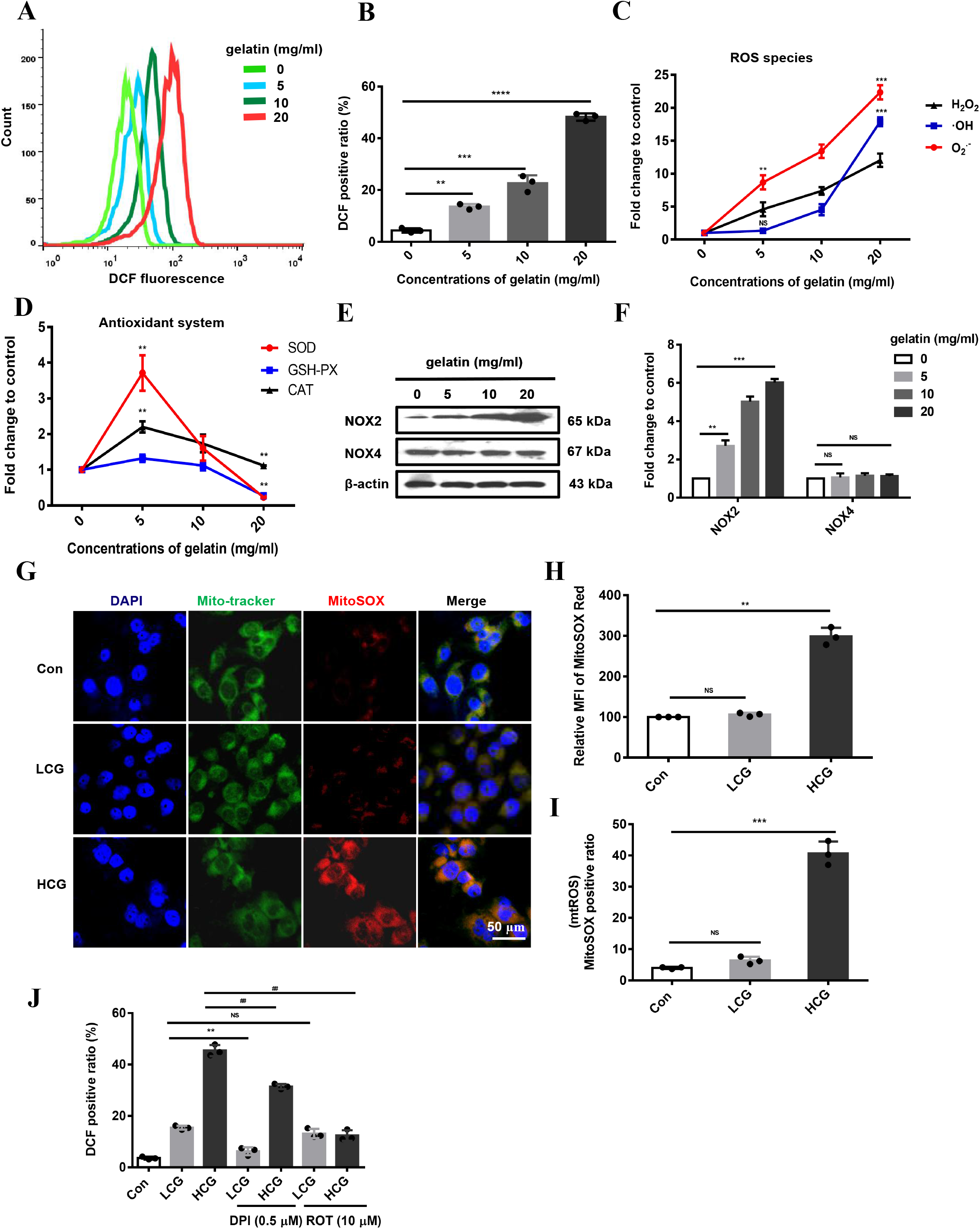
Linear and bell-shaped dose-responses of ROS and antioxidant system to gelatin. **A-B.** Flow-cytometry analysis of cells stained with DCFH-DA (ROS indicator). **C.** Production of specific ROS (O_2_^.−^, ·OH and H_2_O_2_). **D.** Activities of the antioxidases; SOD, GSH-PX and CAT. **E-F.** Western-blots of NOX2 and NOX4 in the cells cultured on gelatin-coated dishes at 90% confluence. β-Actin is used as a loading control. **G-H.** Confocal images of the cells stained with MitoTracker Green, MitoSOX Red dye, and DAPI (blue). Scale bar, 50 μm. **I.** Quantification of MitoSOX-positive cells obtained by flow cytometry. **J.** Flow-cytometry analysis of the cells treated with DPI (0.5 μM, NOX2 inhibitor), and rotenone (ROT 10 μM, inhibitor of mitochondrial respiratory complex I). Significance was determined by unpaired two-tailed Student’s t test analysis. ^##,^ \*\**P* < 0.01; \*\*\**P* < 0.001; \*\*\*\**P* < 0.0001. NS, not significant. n = 3. Data are means ± SEM.

Then we sought to identify the cellular source for gelatin-induced ROS production. ROS can be produced by multiple sources including NADPH oxidases (NOXs) and mitochondrial electron transport complex. NOXs are a vital enzyme family to catalyze ROS production, of which NOX2 and NOX4 are the main isoforms found in skeletal muscle. It is known that NOXs participate in skeletal muscle metabolism and function through redox regulation [30]. Western-blots showed that expression of NOX2 but not NOX4 was upregulated with gelatin in a dose-dependent manner (Fig. 4E-F). We next examined O_2_^.−^ and OH· produced in mitochondria, another significant source of ROS production from oxygen consumption. MitoSOX Red, a selective fluorescence dye for mitochondrial ROS, was used to stain C2C12 followed by imaging (Fig. 4G-H) and flow-cytometry analysis (Fig. 4I). Results show that HCG specifically promoted the generation of mitochondrial ROS, indicating that multiple cellular sources of ROS production can be induced by gelatin. To verify the above findings, we applied specific inhibitors of NOX2, and of complex I of the mitochondrial respiratory chain, diphenyliodonium (DPI) and rotenone (ROT), respectively to C2C12 cultured on LCG and HCG-coated dishes. DCFH-DA staining and flow-cytometry analysis showed that DPI (0.5 μM) treatment suppressed ROS production in both LCG and HCG group, whereras ROT (10 μM) treatment specifically blocked ROS overproduction by HCG (Fig. 4J). Together with ROS species measurements (Fig. 4C), these results indicate that LCG primarily induces production of O_2_^.−^ by NOX2; HCG not only further upregulates NOX2, but also stimulates the production of O_2_^.−^ and OH·by mitochondrial respiratory complex I. The markedly high level of ROS under HCG condition may also be partially attributed to the inhibition of antioxidant activities of SOD and GSH-PX.

### Hormetic function of ROS in myoblast cellular behaviors

To understand the function of ROS on myoblast behaviors, different concentrations of *t*BHP, a donor of O_2_^.−^ and OH·, were added to the culture medium. Previous study revealed that low dose of *t*BHP (lower than 25 μM) moderately promoted the levels of ROS, enhancing proliferation and migration of 3T3-L1 fibroblast cells, while high concentration of *t*BHP (50 μM) caused detrimental accumulation of ROS that inhibited cell behaviors [28]. In this study, we added *t*BHP (5, 10, 50 μM) to C2C12 culture, and confirmed dose-dependent induction of ROS using flow-cytometry analysis (Supplementary Fig. 3A). At 10 μM, *t*BHP promoted growth, migration, and myogenic differentiation of C2C12 cells, but at 50 μM, *t*BHP had opposite effects in these cellular behaviors, suggesting a hormetic effect of ROS (Supplementary Fig. 3B-F). Next, NAC, a scavenger of ROS, was added to the cultures. Removal of ROS was confirmed using flow cytometry (Supplementary Fig. 4A). NAC treatment decreased the proliferation and migration of cells cultured on LCG and reversed the inhibitory effect of HCG on cell behaviors (Fig. 5A-B). NAC reduced both the beneficial effect of LCG and the deleterious effect of HCG in regulating genesis of MyHC-positive cells and fusion of myotubes (Fig. 5C-E) (Supplementary Fig. 4B-C). The results were further supported by the expression levels of mRNAs and proteins of myogenic factors, MyoD, myogenin and MyHC, (Supplementary Fig. 4B-D), thus directly demonstrating a mechanism of hormesis through ROS generation that underlies gelatin’s effect on myogenesis.

**Fig. 5.**
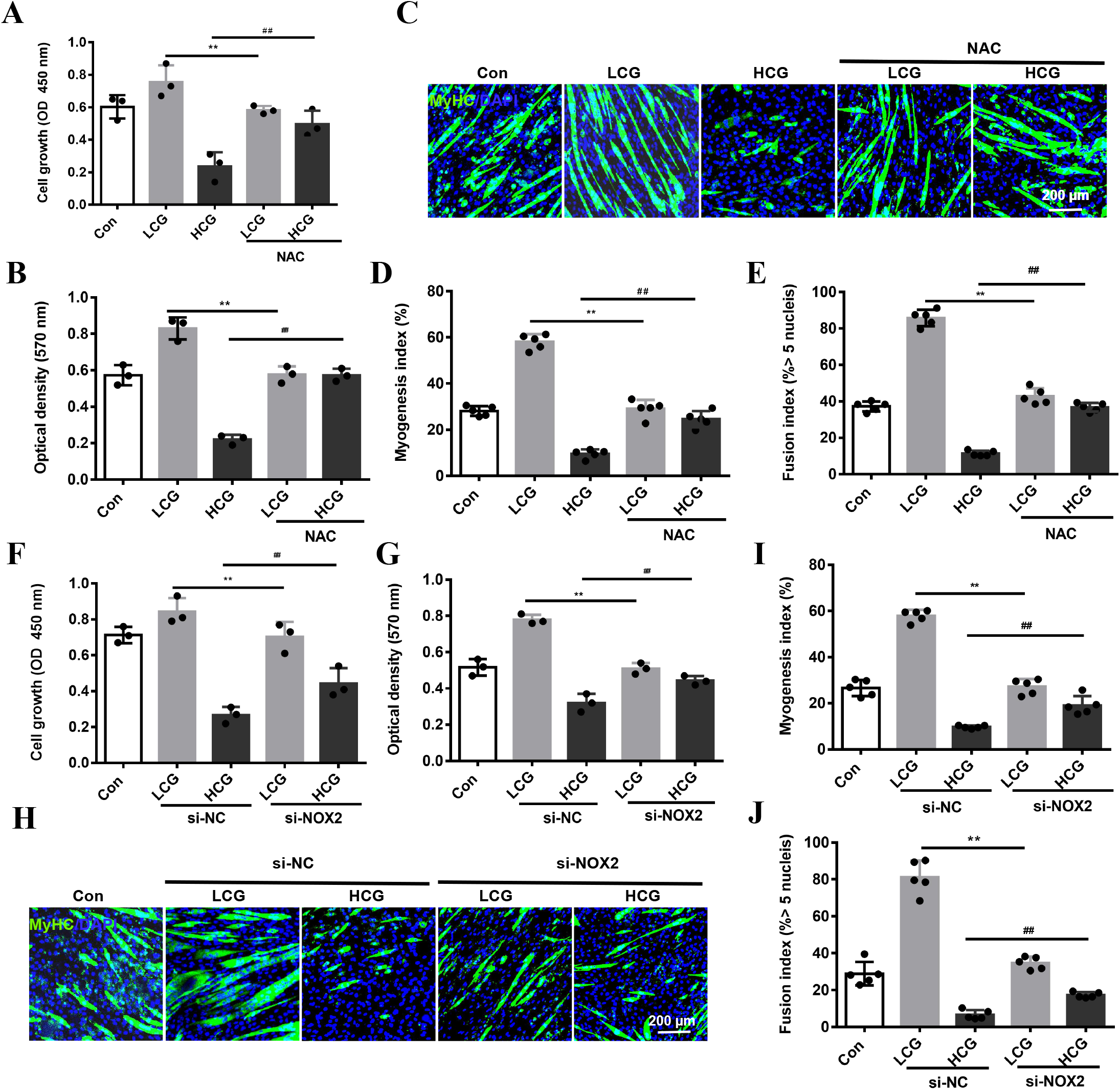
ROS signaling is required for the bi-phasic effect of gelatin. **A-B.** The growth and migration of C2C12 treated with NAC, an ROS scavenger (8 mM). **C.** Confocal images of myogenic differentiation of cells stained with MyHC antibody. Scale bar, 200 μm. **D-E.** Myotube fusion index and myogenesis index were calculated using Image-Pro Plus software. **F-G.** Growth and migration of cells after silencing NOX2 expression (siNOX, or siNC as negative control). **H.** Confocal images of myogenic differentiation of cells stained with MyHC antibody. Scale bar, 200 μm. **I-J.** Myotube fusion index and myogenesis index were calculated using Image-Pro Plus software. Significance was determined by unpaired two-tailed Student’s t-test analysis. ^##, **^*P* < 0.01. n = 3. Data are means ± SEM.

Finally, to verify the role of NOX2 in gelatin effect, we knocked down NOX2 using siRNA, and confirmed reduced expression of NOX2 and lowered levels of ROS production in LCG and HCG (Supplementary Fig. 4G-H). NOX2 silenced inhibition of the growth (Fig. 5F) and migration (Fig. 5G) of the cells on LCG, while it alleviated the inhibitory effect of HCG on cell behaviors. Down-regulation of NOX2 expression also decreased the higher number of MyHC-positive cells and the formation of myotubes (Fig. 5H-J) with a narrower diameter, a shorter length compared with cells on LCG (Supplementary Fig. 4I-J). By contrast, the transfection of NOX2 siRNA apparently increased the myogenic differentiation of cells on HCG (20 mg/ml) (Fig. 5H-J). These data indicated that NOX2 is an important upstream factor in myoblast to mediate gelatin effect.

### IL-6 and TNFα mediate hormetic ROS function for regulating myogenesis

Myokines produced and released by skeletal muscle influence important biological functions of myocytes. They are known to be induced by ROS during physical exercise and muscle regeneration [31]. To identify specific myokines that mediate gelatin and ROS effect, we used ELISA kits to screen potential targets and identified IL-6, whose secretion was promoted by LCG but slightly inhibited by HCG (Fig. 6A). In contrast with IL-6, the release of TNFα was specifically induced by HCG (Fig. 6B). Other myokines such as pro-inflammation factors, IL-1β and IL-18 as well as chemokines, CCL5 and CCL2/MCP-1, were unaffected by different doses of gelatin (Supplementary Fig. 5A-D). To determine the role of ROS in myokine release, we added ROS donor *t*BHP to C2C12 cultures at different concentrations, which induced release of IL-6 and TNFα in a parallel manner with gelatin (Supplementary Fig. 5E). Furthermore, treatment of NAC (ROS scavenger) specifically blocked release of IL-6 from LCG myoblast and TNFα from HCG myoblast, indicating ROS are responsible for the activation of both myokine pathways (Fig. 6C).

**Fig. 6.**
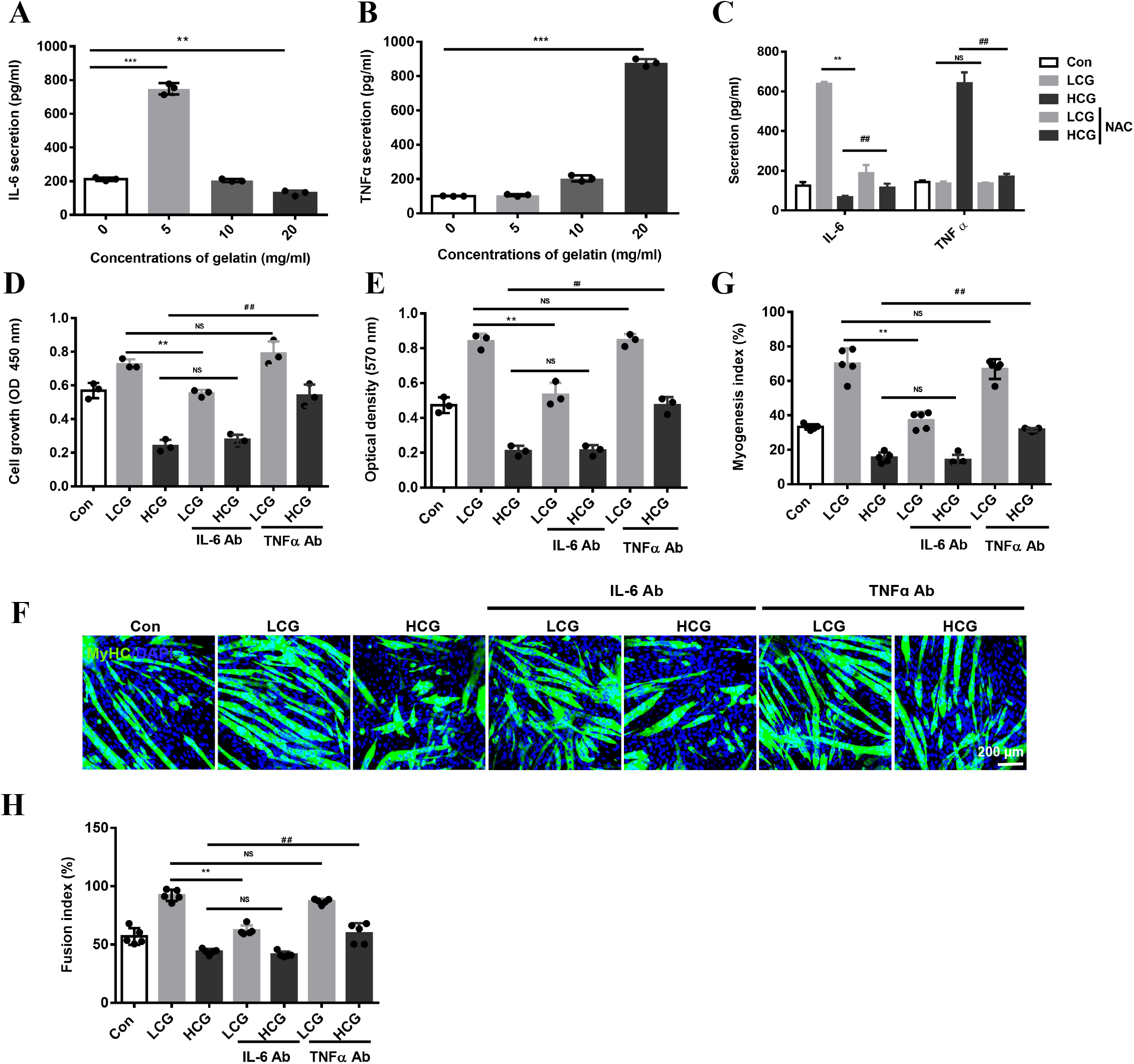
IL-6 and TNFα mediate dual regulation of ROS on myogenesis. **A-B.** ELISA analysis of IL-6 and TNFα in the culture supernatants of C2C12 cells on gelatin. **C.** ELISA analysis of IL-6 and TNFα in the culture supernatants of C2C12 cells treated with NAC. **D-E.** Growth and migration of the C2C12 myoblasts incubated with the neutralizing antibodies (Ab) to IL-6 (0.3 μg/ml) and TNFα (0.5 μg/ml). **F.** Confocal images of myogenic differentiation of cells stained with anti-MyHC antibody after incubation with IL-6 and TNFα. Scale bar, 200 μm. **G-H.** The myogenesis index and myotube fusion index were calculated using Image-Pro Plus software. Significance was determined by unpaired two-tailed Student’s t-test analysis. ^##, **^*P* < 0.01; ^***^*P* < 0.001. NS, not significant. n = 3. Data are means ± SEM.

To directly confirm the function of IL-6 and TNFα in gelatin-induced myogenesis, purified mouse recombinant myokines were exogenously added to C2C12 cultures. The results showed that IL-6 enhanced the growth and migration of cells and increased the number of MyHC-positive cells with a wider diameter and a longer length compared with control, whereas TNFα exerted an opposite effect (Supplementary Fig. 6A-G). Then we investigated whether the release of IL-6 and TNFα is necessary for the gelatin effect by the addition of IL-6 and TNFα neutralizing monoclonal antibodies (Ab) in the culture medium. Enhanced growth and migration of cells on LCG were decreased by IL-6 Ab but not by TNFα Ab (Fig. 6D-E). By contrast, the inhibitory effect by HCG was reversed by TNFα Ab but not by IL-6 Ab (Fig. 6D-E). The blockade of IL-6 effectively decreased the number of MyHC-positive cells and inhibited the cell fusion (Fig. 6F-H), which resulted in the formation of myotubes with a narrower diameter and a shorter length compared with cells on LCG (Supplementary Fig. 6H-I). By contrast, anti-TNFα Ab had no effects on LCG cells, but significantly reversed the HCG effect on myogenesis and expression of myogenesis-related proteins, MyoD, myogenin and MyHC (Supplementary Fig. 6J-K). These data reveal that the bi-phasic effect of gelatin on myogenesis can be mediated by IL-6 and TNFα that are induced by low and high levels of ROS, respectively.

### Gelatin-induced myogenesis in mouse primary myoblasts

In order to verify the gelatin-ROS-IL-6/TNFα signaling cascade in myogenesis that we have identified in C2C12 cells, we repeated experiments in mouse primary myoblasts (MPMs) isolated from TA muscle [32]. The primary TA muscle cells adhered to the plates with a round morphology within 4 hours after plating and grew into flat fibroblast-like morphology, with desmin (myoblast marker) detectable in more than 95% cell population (Supplementary Fig. 7A-B). After reaching 90% confluence, cells were subjected to 2% horse serum to induce myogenic differentiation (Supplementary Fig. 7A). Consistent with the results in C2C12 cells, the growth and migration of MPMs were enhanced by LCG but inhibited by HCG (Fig. 7A-B). LCG increased MyHC-positive cells (Fig. 7C-D) and promoted the formation of myotube with a wider diameter and a longer length compared with control, while HCG exerted opposite effects (Supplementary Fig. 7C-E). ROS production and the NOX2 expression in MPMs was induced by gelatin in a similar manner to C2C12 (Fig. 7E-F). Removal of ROS by NAC (Supplementary Fig. 7F) not only decreased the LCG-enhanced growth and migration of cells but also reversed the inhibitory effect of HCG on cell proliferation and migration (Fig. 7H-I). NAC also decreased the MyHC-positive cells (Fig. 7J-K), myotube formation, the length and width of myotube in LCG group, while increasing myogenesis in HCG cells (Supplementary Fig. 7G-I). LCG significantly promoted IL-6. HCG slightly inhibited IL-6 but significantly enhanced TNFα (Fig. 7G). The suppression of ROS by NAC reduced IL-6 and TNFα in culture medium in TA muscle myoblasts cultured on LCG and HCG substrates (Fig. 7l). These data suggested that the signaling cascade of gelatin-ROS-IL-6/TNFα underlying the bi-phasic effect of gelatin is valid in primary muscle cells.

**Fig. 7.**
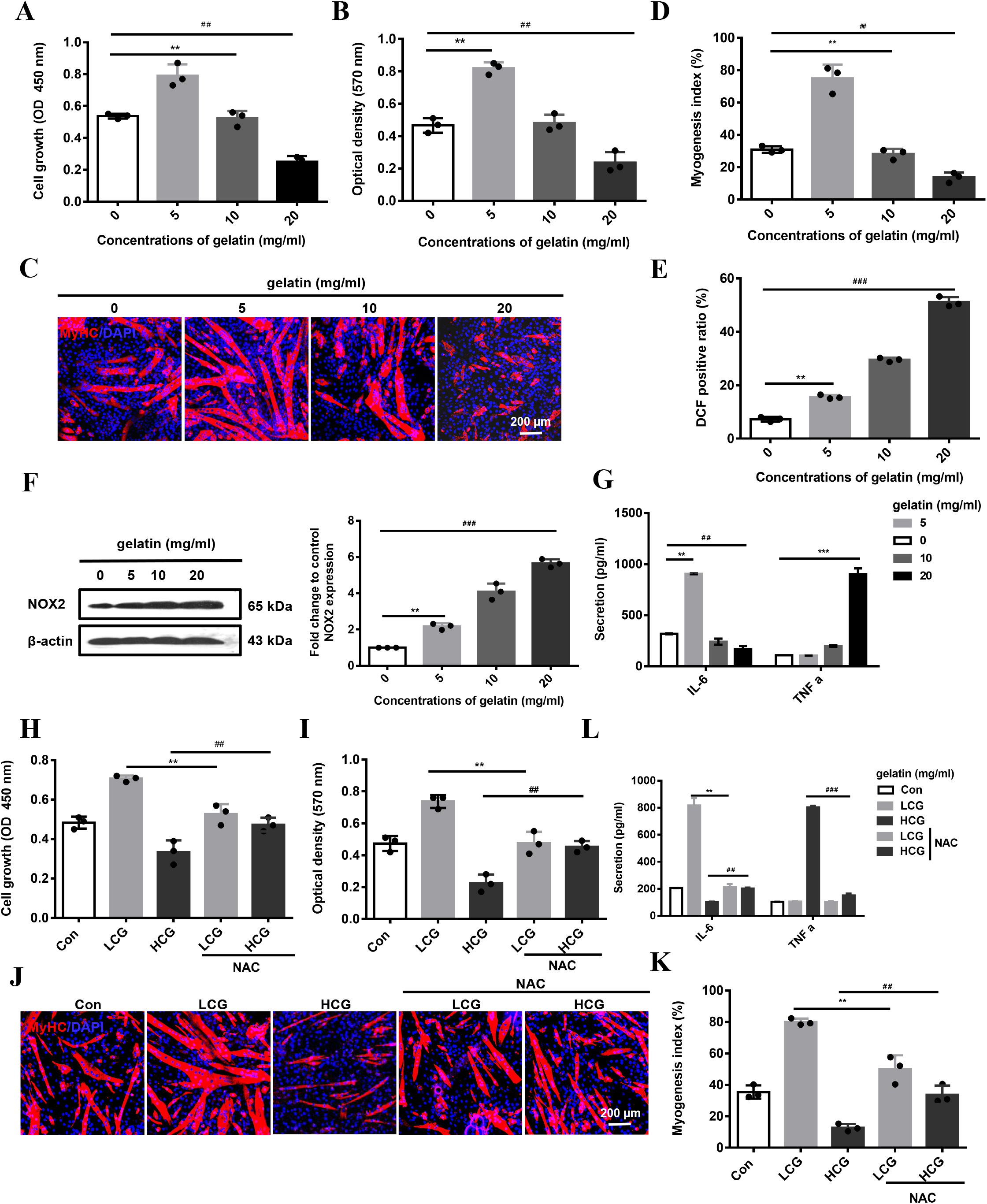
ROS and cytokines signaling mediate bi-phasic effect of gelatin substrate in myogenesis of mouse primary myoblasts (MPMs). **A-B.** Growth and migration of MPMs show bell-shaped concentration-response to gelatin substrate. **C.** Confocal images of differentiated MPMs stained with anti-MyHC antibody. Scale bar, 200 μm. **D.** The myogenesis index calculated using Image-Pro Plus software. **E.** Flow-cytometry analysis of MPMs stained with DCFH-DA. **F.** Western-blots of NOX2 proteins in the MPMs cultured on increasing concentrations of gelatin substrate. β-Actin is used as a loading control. **G.** ELISA analysis of IL-6 and TNFα levels in the culture supernatant of MPMs. **H-I.** Growth and migration of the MPMs treated with NAC, an ROS scavenger (8 mM). **J.** Confocal images of differentiated MPMs stained with anti-MyHC antibody. Scale bar, 200 μm. **K.** The myogenesis index was calculated using Image-Pro Plus software. **L.** ELISA-analysis of the levels of IL-6 and TNFα in the culture supernatant of MPMs treated with NAC (8 mM). Significance was determined by unpaired two-tailed Student’s t-test analysis. ^##, **^*P* < 0.01; ^###, ***^*P* < 0.001. n = 3. Data are means ± SEM.

### Gelatin-ROS-IL-6/TNFα cascade underlies skeletal muscle regeneration in vivo

Finally, we examined the oxidative situation in the gelatin-injected TA muscle at 14 D.P.I. DCFH-DA staining showed that ROS level was upregulated modestly by LCG as well as the antioxidation activities of SOD and GSH-PX. Furthermore, high accumulation of ROS and reduced activities of SOD and GSH-PX were observed in HCG animals (Fig. 8A-D). The overaccumulation of ROS by HCG injection also caused a higher level of malondialdehyde (MDA) detected in the regenerated muscle, indicating higher oxidative stress, whereas LCG marginally reduced MDA (Fig. 8E). We further examined NOX2 expression and found that the protein was not only upregulated during natural muscle regeneration but was further increased by gelatin injection in a dose-dependent manner (Fig. 8F-I). Lastly, we examined IL-6 and TNFα in 14 D.P.I. TA muscles. The results showed an increase of IL-6 cytokine in the regenerating muscle which was further enhanced by LCG but slightly inhibited by HCG. By contrast, dramatically induced TNFα was detected specifically in HCG animals (Fig. 8J).

**Fig. 8.**
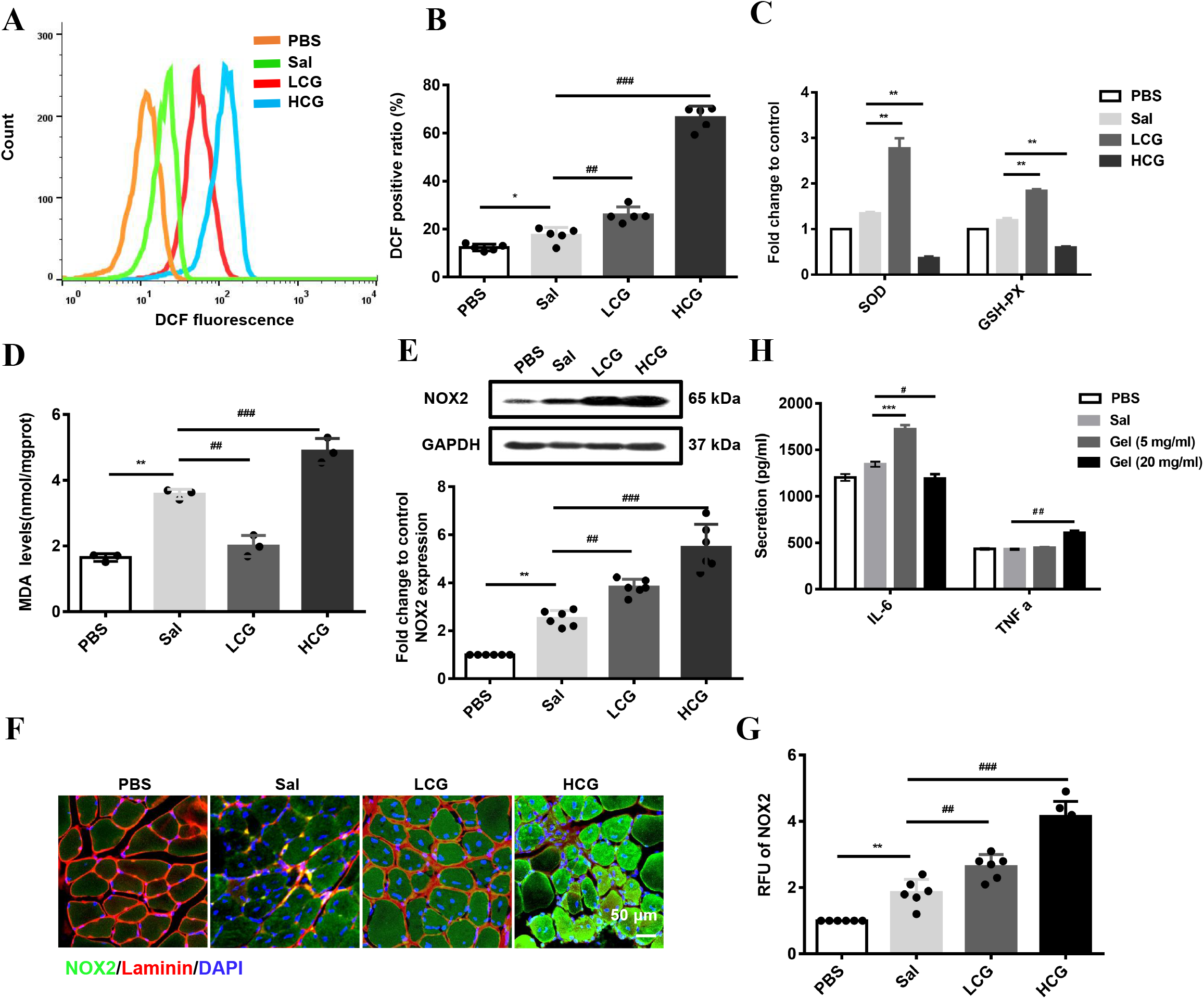
ROS, antioxidant, and cytokine signaling cascade upon low- and high-dose gelatin injection during muscle regeneration. **A-B.** Flow-cytometry analysis of ROS in damaged TA muscle cells stained with DCFH-DA (n=5). **C-D.** The activities of SOD and GSH-PX in damaged TA muscle (n=3). **E.** MDA level in the damaged TA muscle cells (n=3). **F-G.** Western-blots of NOX2 in the damaged TA muscle (n=6, GAPDH as a loading control). **H-I.** IHC analysis of NOX2 (green) in the damaged TA muscles stained with laminin (red) and DAPI (blue) at 7 D.P.I (n=6). The relative fluorescence unit of NOX2 myofibers per filed is shown on the right. **J.** ELISA-analysis of IL-6 and TNFα in damaged TA muscle tissue at 7 D.P.I (n=3). Significance was determined by unpaired two-tailed Student’s t-test analysis. ^##, **^*P* < 0.01; ^###, ***^*P* < 0.001. Data are means ± SEM.

In summary, the novel gelatin-ROS-IL-6/TNFα cascade during skeletal muscle regeneration found in the present study is illustrated (Fig. 9).

**Fig. 9.**
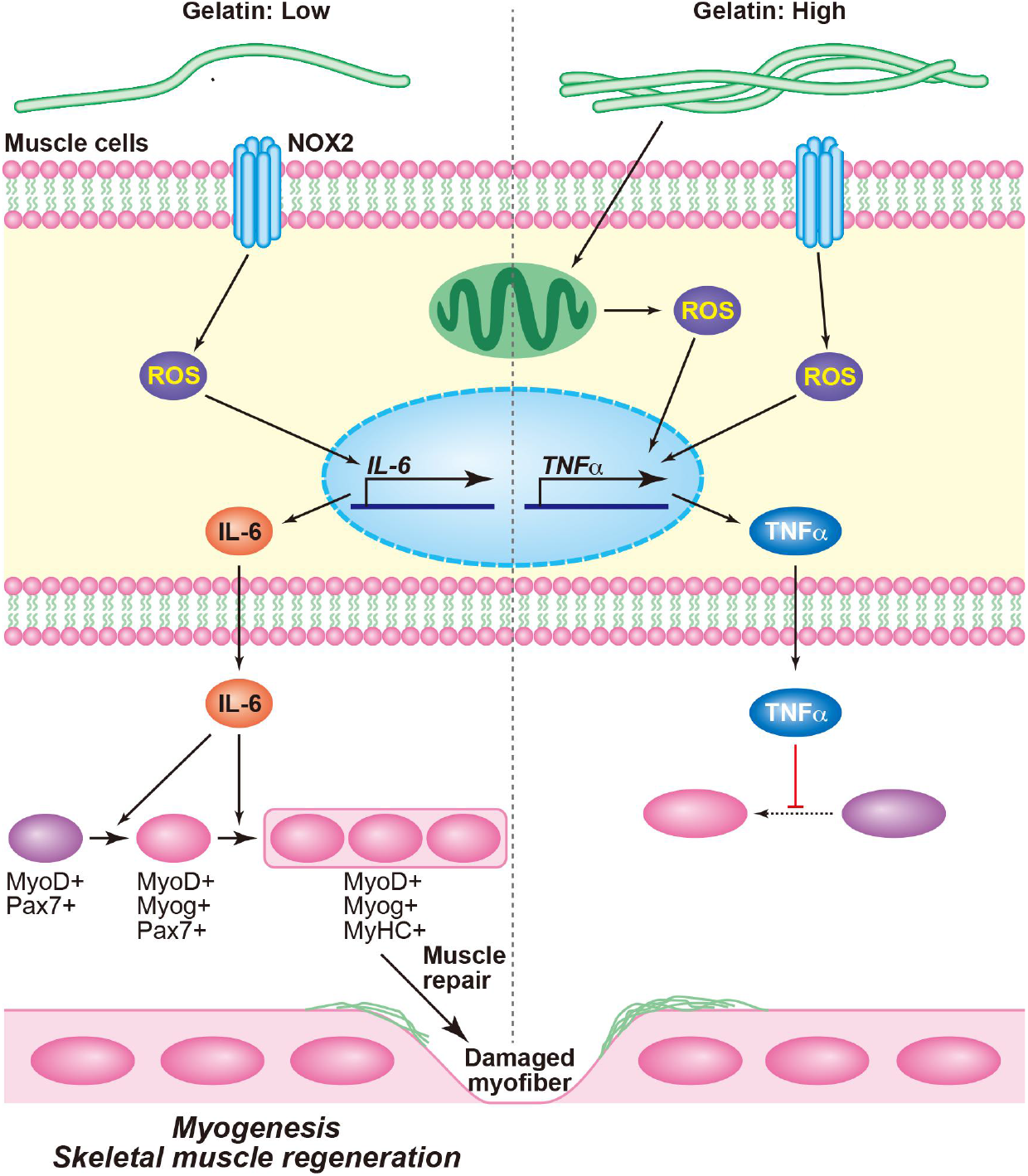
ROS-antioxidant-cytokine signaling cascade in mediating bi-phasic effect of gelatin in myogenesis and skeletal muscle regeneration. Low-dose gelatin promotes myogenesis and muscle repair via moderate production of ROS and antioxidant generation, inducing IL-6 secretion. High-dose gelatin hampers myogenesis and muscle regeneration by triggering overproduction of ROS by NOX2 and mitochondria as well as by inducing TNFα secretion.

## Discussion

We have discovered that a modest injection of gelatin into injured skeletal muscle can aid the regeneration process. We also found pleiotropic effects of gelatin on myogenesis in vitro and in vivo mediated by intracellular ROS-IL-6/TNFα signal cascade, which is reminiscent of hormesis in toxicology. These results suggest that temporary production and accumulation of gelatin during ECM remodeling in muscle regeneration facilitates injury recovery.

Gelatin is the natural product of collagen during ECM remodeling in skeletal muscle regeneration [33]. Although a large amount of studies focused on the application of gelatin-based materials in muscle engineering, it was unclear whether gelatin influences the biological function of myoblasts and muscle regeneration. Our study has revealed complex effects of gelatin on myogenesis in vitro and skeletal muscle regeneration in vivo through ROS signaling pathway. The apparently paradoxical findings of gelatin can be explained within the framework of hormesis, defined as a biphasic dose-response. The present work firstly illustrates the occurrence of hormesis in biological function of ECM proteins and further proven hormesis appears to be independent of biological model; in vitro cultured cells or in vivo. The therapeutic or toxic effect of gelatin on skeletal muscle regeneration is changeable in a dose-dependent manner. LCG plays a beneficial role in muscle regeneration, but HCG aggravates the muscle damage to delay repair. The adverse effect of HCG on myogenesis might be attributed to the aging or growth arrest of cells but the specific mechanism of HCG-caused damage is needed for further investigation. The safety profile of gelatin imposes a careful analysis of the risk/benefit balance prior to proposing clinical application.

As the hydrolysate of collagen, gelatin (non-fibrous without triple-helicity) is locally and temporally produced in skeletal muscle during inflammation and regeneration [33]. It was reported that gelatin fibers artificially produced by spinning induced cell aggregation in rabbit skeletal muscle myoblasts and regulated aligned muscle tissue formation [33]. Gelatin hydrogels promotes the myogenesis of human muscle progenitor cells [34]. This effect may be mediated by RGD (R, arginine; G, glycine; D, aspartate) domain interacting with integrin receptors (αvβ3, α5β1 and α7β1) at cell surface [35]. It was shown that gelatin coating facilitates adhesion and spread of C2C12 myoblast through integrin receptors (αvβ3 and α5β1) [36]. Integrin receptors influence F-actin assembly to regulate the cell morphological changes through FAK/Rho A/ROCK pathway [37]. In this study, gelatin induced the changes of cell morphology and migration, suggesting that integrin receptors might be the mediator of gelatin.

There is growing evidence supporting the effect of gelatin on ROS generation and redox function. Gelatin enhances ROS to induce the aggregation of peritoneal macrophages and release of pro-inflammatory factors [38]. Gelatin facilitates the production of myofibroblasts from 3T3-L1 and C2C12 cells via ROS. Ma et al. reported that gelatin hydrolysates can protect the mouse embryonic fibroblasts from UVB irradiation via ROS pathway [39]. Gelatin-derived peptides also possess antioxidant properties through multiple pathways such as inhibiting lipid peroxidation, scavenging free radical, and chelating transition metal ion [39]. Peptides isolated from fish skin gelatin were shown to protect liver cells from oxidative damage by ROS donor *t*BHP [40]. These studies have indicated that gelatin exert pleiotropic functions through ROS signal in different tissues.

We found that gelatin triggers ROS production both by the NADPH oxidase, NOX2, and mitochondrial respiratory complex I. Little is known about the function of gelatin on NADPH system, except for one report, showing that gelatin gel might increase the resistance of NADPH-oxidoreductase enzyme system [41]. Collagen is shown to induce O_2_^.−^ via regulating the activity of NOX at p47 (phox) (the organizer subunit) during platelet activation [42, 43], but how gelatin affects NADPH system remains to be elucidated. Similar to HCG, gelatin nanoparticles have been reported to decrease activities of SOD, GSH-PX and CAT during apoptosis of NCI-H460 lung cancer cells [44]. Although gelatin influences the generation of ROS in different biological processes, underlying mechanisms are not fully understood.

Skeletal muscle is an endocrine tissue as it produces and secretes cytokines, also known as myokines, growth factors, and pro-inflammatory factors, which play important roles in muscle regeneration [45]. ROS are related to the induction and accumulation of myokines such as IL-6 in myogenesis [46]. IL-6 is a multifunctional cytokine not only as a pro-inflammatory factor, but also as a myokine released from muscle in exercise or regeneration [47]. It has been reported that the level of IL-6 in muscle can be transiently elevated to 100-fold after exercise for 4 h [48]. IL-6 exerts a critical role on muscle homeostasis and diseases, and is related to the hypertrophic muscle growth and myogenesis via regulating muscle stem cells [49, 50]. Our finding in facilitated expansion of Pax7^+^ cells by LCG is consistent with this finding. TNFα is a major pro-inflammatory cytokine that is increasingly expressed in damaged or dystrophic muscle [51]. It was shown that high level of TNFα may cause apoptosis of myoblasts and myocytes, thus causing muscle atrophy [52]. Previous studies have shown that collagen I influences the migration and differentiation of myoblasts through regulating release of IL-6 [9]. Gelatin has also been shown to regulate production and secretion of IL-6 and TNFα in differentiated U937 lymphoma cells [18]. In the present report, we show that gelatin can induce IL-6 and TNFα in myoblasts at different concentrations.

In addition to TNFα, gelatin has also been shown to induce IL-1β and prostaglandin E_2_ in macrophages [38]. *Kojima* et al. reported that bovine bone gelatin stimulates the secretion of cytokines IL-6, MCP-1 and MIP-2 in murine adherent spleen cells to regulate cell proliferation [53]. However, how gelatin and ROS functions diverge through IL-6 and TNFα signalings in myoblasts remains unclear. One possibility is that the different amount and species of ROS lead to activation of specific pathways. ROS and myokines have both be shown to promote muscle adaptation to exercise [31]. It was reported that IL-6/STAT3 pathway participate in myoblast proliferation and macrophage migration during muscle regeneration [54]. TNFα/NF-κB p65 pathway inhibits the myogenic differentiation of myoblasts, implicates as a mediator of muscle wasting [55]. The versatile downstream myokines may further diversify cell adaptation behaviors to ECM and gelatin, warranting further investigation.

Detailed investigation of interplay between ECM remodeling and intracellular signaling pathways for skeletal muscle regeneration has yet to be conducted. Given the physiological importance and promises of using multifunctional gelatin as a biomaterial in tissue engineering and regenerative medicine, better elucidation of such fundamental pathways is likely to give rise to therapeutic value.

## Materials and Methods

### Animals

All experiments were performed according to P. R. China legislation on the use and care of laboratory animals and the criterion confirmed by the Institute for Experimental Animals at Shenyang Pharmaceutical University. C57/BL6 mice at the age of 6-8 weeks, weighting 18-22 g, were obtained from Changsheng Biotechnology (Liaoning, China). Mice were maintained in light-(12 h light/12 h dark cycle), climate-(temperature 23 ± 0.5 °C, humidity 55-65%) controlled environment and fed a normal chow diet with water *ad libitum*.

### Skeletal muscle injury and gelatin injection

10-weeks old C57/BL6 male mice were anesthetized and injected with PBS or cardiotoxin (CTX, 20 μL, 10 μM) in hindlimb TA muscle. At 2 d postinjury (2 D.P.I.), mice were injected with either saline or gelatin (5 and 20 mg/ml in saline) into injured TA muscle. The muscles were harvested at 7 and 14 D.P.I. for the analyses.

### Histological analysis

TA muscles were dissected from mice, embedded into OCT compound and quick-frozen by immersing in liquid nitrogen, sectioned on a cryostat (AS-620; Shandon, Astmoor, UK) at a thickness of 10 μm. The sections were further stained with hematoxylin and eosin (H&E) that was conducted on 10% formalin-fixed muscle section with Mayer’s H&E (Sigma Chemical, St. Louis, MO, USA). Immunofluorescence (IHC) staining was also performed for subsequent further analysis.

### Reagents

Primary antibodies against MyoD (18943-1-AP) and myogenin (67082-1-Ig) were purchased from Proteintech (Wuhan, Hubei, China). Primary antibodies against β-actin (BF0198), GAPDH (AF7021), Pax7 (AF7584), desmin (AF5334), NOX2 (DF6520) and NOX4 (DF6924) were obtained from Affinity Biosciences (Affinity Biosciences. OH. USA). MyHC (MAB4470), monoclonal neutralization antibodies against IL-6 (AB-406-NA) and TNFα (AB-410-NA), and mouse recombinant proteins of IL-6 and TNFα were brought from R&D system (Minneapolis, MN, USA). Mouse anti-eMyHC was purchased form Developmental Studies Hybridoma (DSHB, Iowa, IA, USA). Fluorescein (FITC)-conjugated AffiniPure goat anti-mouse IgG (H+L) was obtained from KeyGEN Biotechnology (Nanjing, Jiangsu, China). Rabbit anti-laminin antibody (cat. no. L9393), 2′,7′-dichlorodihydro fluorescein diacetate (DCFH-DA), N-acetyl-L-cysteine (NAC), tertiary butylhydroperoxide (*t*BHP), diphenyliodonium (DPI) and rotenone (ROT) were purchased from Sigma Chemical (St. Louis, MO, USA).

### Isolation and differentiation of MPMs

Primary myoblasts were isolated from 3-day-old C57/BL6 mice. The hindlimb muscles were aseptically dissected, minced mechanically and digested with 0.2% collagenase II (Invitrogen, cat. 17101015, Carlsbad, CA, USA) and 0.05% trypsin in PBS for 40 min at 37 °C with slight agitation. Then the mixture was filtered with a 200 mesh (75 μm) filter and centrifuged at 400 x g for 5 min. Cells were resuspended in PBS, and separated in 20%/60% percoll (Solarbio, 65455-52-9, Beijing, China) by centrifugation at 800 x g for 20 min. Then the cells were resuspended in growth Ham’s F10 medium (Gibico, 11550043, NY, USA) containing 20% FBS (Gibico, 10099141, NY, USA), 2.5 ng/ml basic FGF (Cyagen, HEGFP-0602, Suzhou, China) and 1% penicillin/streptomycin and were cultured on plates (Corning, Corning, NY, USA) for 2 h; the non-adherent cells were then transferred to another dish. For inducing myogenic differentiation, culture medium of cells at 90% confluence was replaced with DMEM containing 2% horse serum (26050070, Gibco, NY, USA). Cells were cultured in the differentiation medium for another 5 days until formation of myotubes.

### Cell culture and myogenic differentiation

C2C12 myoblast cells (American Type Culture Collection, Manassas, VA, USA) were cultured in DMEM (high glucose, Gibco, NY, USA) supplemented with 10% fetal bovine serum (FBS) (Beijing Yuanheng Shengma Research Institution of Biotechnology, Beijing, China), 100 μg/ml streptomycin and 100 U/ml penicillin, at 37 °C and 5% CO_2_. For inducing myogenic differentiation, culture medium of cells at 90% confluence was replaced with DMEM containing 2% horse serum (Gibco, NY, USA). Cells were cultured in the new medium for further 5 days until the formation of myotubes.

### Preparation of gelatin-coated dishes

Gelatin was prepared in Nippi Research Institute of Biomatrix (Toride, Ibaraki, Japan) and was diluted in 0.5 mM acetic acid to the indicated concentrations (5, 10, 20 mg/ml) and added to cell culture dishes (Corning Inc., Coring, NY), incubated for 4 h at 37 °C, 5% CO_2_. Before plating cells, gelatin solution was discarded and the remaining was rinsed off by PBS three times [17].

### Cell growth and migration detection

Cell growth was examined using a CCK-8 kit (Beyotime Biotechnology, Shanghai, China) according to manufacturer’s instruction. Migrations were determined by cell-scratch and transwell cell migration assays as described previously [9][26].

### Confocal fluorescence microscopic analysis

The indicated cells were fixed with paraformaldehyde (PFA) for 20 min and permeabilized with 0.15% Triton X-100 for 12 min. After that, the cells were blocked with PBS containing 10% FBS for 30 min. For the cell morphology analysis, the cells were incubated with Alexa Fluor 488 Phalloidin (1:500, Ye Sen, Shanghai, China) overnight at 4 °C. For MyHC and desmin staining, cells were incubated with anti-MyHC antibody (1:50) or desmin (1:100) overnight at 4 °C in PBS, and then treated with fluorescent secondary antibody (1:200) for 2 hours at RT in dark. After the incubation with secondary antibody, cells were stained with DAPI (1 μg/ml) (Beyotime Biotechnology, Shanghai, China) for 10 min, and imaged with a confocal microscope (Nikon, Tokyo, Japan).

### Immunohistochemistry

TA muscle sections were fixed with 4% PFA and permeabilized with 0.2% Triton X-100 for 10 min in PBS. After that, sections were incubated with 3% H_2_O_2_ for 20 min to remove the endogenous peroxidase and blocked with 10% normal goat serum for 2 h at room temperature. Sections were incubated with Alexa Fluor 488 Phalloidin (1:300) and primary antibody against eMyHC (DSHB, 1:50), laminin (1:100), Pax7 (1:300) and MyoG (1:300) at 4 °C overnight. Last, sections were treated with FITC- or TRITC-conjugated secondary antibodies for 2 h and DAPI for 10 min to observe the muscle with a confocal microscope (Nikon, Tokyo, Japan).

### 5-Ethynyl-2-deoxyuridine (EdU) staining assay

Cells cultured on gelatin-coated dishes for 24 h were incubated with EdU (10 μM) (C0071S, Beyotime Biotechnology, Shanghai, China) for 2 h, fixed in 4% PFA, permeabilized by 0.5% Triton X-100, and then detected according to manufacturer’s protocol with hoechst 33342 for cell nuclei.

### ELISA analysis

Cells (3.0×10 ^5^ cells/well) were cultured on gelatin-coated dishes until reaching 90% confluence. Then cells and supernatants were harvested separately, and the cell number in each well was counted under a microscope. The amount of detected myokines was normalized to cell number in each well. In muscles, TA muscles were minced by a grinder and crushed by ultrasonication in PBS (1:9). The homogenate was centrifuged at 800 x g for 20 min and supernatants were collected. Cytokines, IL-6, CCL2/MCP-1, CCL5, TNFα, IL-1β and IL-18, in cell cultures and IL-6 and TNFα in muscle tissue were examined using ELISA kits (DakeweBiotech, Shenzhen, China) following the manufacturer’s protocols.

### Gene expression analysis

Total RNA was extracted using RNAiso plus reagent (TaKaRa, Tokyo, Japan) following manufacturer’s instructions. 1 μg RNA was used to synthesize first strand of cDNA using PrimeScriptTM RT-PCR Kit (TaKaRa). qPCR was conducted using PrimeScriptTMRT Master Mix (TaKaRa) on ABI 7500 Fast to quantify mRNA of MyoD, myogenin and MyHC. Primer sequences are as follows (5’-3’):

MyoD:

forward: CATTCCAACCCACAGAACCT;
reverse: CAAGCCCTGAGAGTCGTCTT;
myogenin:

forward: CAATGCACTGGAGTTCGGT;
reverse: GCCAGGTTGACATTGGATTG;
MyHC:

forward: CGCAAGAATGTTCTCAGGCT;
reverse: GCCAGGTTGACATTGGATT;
GAPDH:

forward: TCCCACTCTTCCACCTTC;
reverse: CTGTAGCCGTATTCATTGTC.

Relative expression of target genes was determined by comparing to GAPDH mRNA. The mRNA index of myogenic proteins was calculated and plotted as an integrated fold change of MyoD, myogenin and MyHC mRNA to control by using scale function in R-package.

### Transfection of siRNA

Cells were transfected with siRNA targeting NOX2 and negative control using Lipofectamin 2000 (Invitrogen, CarIsbad, CA, USA) according to the manufacturer’s instructions. Then the cells were incubated for another 48 h before subsequent experiments.

### Western-blot analysis

The cells and supernatants were harvested to acquire protein samples using RIPA lysis buffer (Beyotime, Haimen, China) containing protease inhibitor cocktail (Solarbio, Beijing, China). TA muscles were minced and crushed by ultrasonication in RIPA lysis buffer (1:9). The homogenate was centrifuged and supernatants were collected. Protein concentrations were examined by BCA (WLA004b, Shenyang, China) according to manufacturer’s instructions. The proteins were separated by the 10-12% SDS-PAGE. After electrophoresis, the proteins were transferred to Millipore Immobilon®-P Transfer Membrane (Millipore, Billerica, MA, USA) and blocked in 5% skin milk solution for 2 hours. Then the membranes were incubated with primary antibodies overnight at 4°C. Next day, the horseradish-peroxidase-conjugated secondary antibodies were added to the membranes and incubated for 3 hours, and developed with SuperSignal® West Pico Chemiluminescent Substrate (Thermo Fisher Scientific, Rockford, IL, USA).

### Enzyme activity and MDA assay

The indicated cells and supernatants were collected and lysed in RIPA buffer. TA muscles were minced and crushed by ultrasonication in pre-cooled saline (1:9). The homogenate was centrifuged and supernatants were collected. Protein concentrations in lysates were quantified using BCA kit (Solarbio, Beijing, China). The content of MDA and the enzyme activities of SOD, GSH-PX and CAT were estimated by using kits (Nanjing Jiancheng Bioengineering Institute, Nanjing, China) according to the manufacturer’s instructions.

### Flow-cytometry analysis of ROS

ROS in the total intracellular and TA muscle were examined using DCFH-DA or MitoSox Red staining (Ye Sen, Shanghai, China). For intracellular and mitochondrial ROS, the indicated cells were incubated with 10 μM DCFH-DA or 5 μM MitoSOX Red at 37 °C for 30 min, washed three times with PBS and analysed using flow cytometry. To visualize mitochondrial ROS, the indicated cells were incubated with 100 nM MitoTracker Green FM and 5 μM MitoSOX Red at 37 °C for 30 min (Ye Sen, Shanghai, China). Then the cells were washed three times with PBS and fixed with PFA for 20 min. Then the cells were permeabilized with 0.15% triton X-100 for 12 min and blocked with 10% FBS for 30 min. Lastly, the cells were stained with DAPI for 10 min, and imaged with a confocal microscope (Nikon, Tokyo, Japan). For TA muscle ROS, the tissues were minced, digested with 0.2% collagenase II and 0.05% trypsin for 40 min to obtain single cell suspension. The cells were resuspended with 10 μM DCFH-DA in PBS for 30 min after the centrifugation, followed by washing three times with PBS and analysed using flow cytometry.

### Statistical analysis

All results were obtained from at least three independent experiments. Data are shown as means ± SEM. The data were analyzed using GraphPad software and Student’s *t*-test was used to examine the comparisons between groups. *P*<0.05 is considered as significant.

## Acknowledgement

This work was financially supported by NSFC 31971335, Xingliao Talents Program XLYC1802007, Department of Education of Liaoning Province 1911520092 to Wang DO, Department of Education of Liaoning Province 2020LQN01 to Liu X and “Chunhui Plan” LN2019016 to Yu Q.

## Author contributions

Liu X, Ikejima T, and Wang DO conceived the project and wrote the paper. Liu X planned experiments, analyzed data and wrote the paper. Zu E, Chang X and Wang Z performed experiments. Kamei K, Li X and Yu Q supervised part of the experiments and contributed to interpretation of the results. Hayashi T, Mizuno K, Hattori S and Fujisaki H provided reagents and revised the paper. All authors provided critical feedback and helped shape the research, analysis and manuscript.

## Materials & Correspondence

Requests for materials and inquiries should be addressed to Wang DO.

